# Extracellular vesicle microRNAs contribute to Notch signaling pathway in T-Cell Acute Lymphoblastic Leukemia

**DOI:** 10.1101/2022.07.20.500780

**Authors:** Tommaso Colangelo, Patrizio Panelli, Francesco Mazzarelli, Francesco Tamiro, Valentina Melocchi, Orazio Palumbo, Giovanni Rossi, Vincenzo Giambra, Fabrizio Bianchi

## Abstract

T-cell acute lymphoblastic leukemia (T-ALL) is an aggressive T-cell malignancy characterized by genotypically-defined and phenotypically divergent cell populations, governed by adaptive landscapes. Clonal expansions are associated to genetic and epigenetic events, and modulation of external stimuli that affect the hierarchical structure of subclones and support the dynamics of leukemic subsets. Recently, small extracellular vesicles (sEV) such exosomes were also shown to play a role in leukemia. Here, by coupling miRNome, bulk and single cell transcriptome profiling, we found that T-ALL-secreted sEV contain NOTCH1-dependent microRNAs (EV-miRs), which control oncogenic pathways acting as autocrine stimuli and ultimately promoting the expansion/survival of highly proliferative cell subsets of human T-cell leukemias. Of interest, we found that NOTCH1-dependent EV-miRs mostly comprised members of miR-17-92a cluster and paralogues, which rescued in vitro the proliferation of T-ALL cells blocked by γ-secretase inhibitors (GSI) and regulate network of genes characterizing patients with relapsed/refractory early T-cell progenitor (ETP) ALLs. All these findings suggest that NOTCH1 dependent EV-miRs may sustain the growth/survival of immunophenotypically defined cell populations, altering the cell heterogeneity and the dynamics of T-cell leukemias in response to conventional therapies.

## Background

Acute lymphoblastic leukemia or ALL, is an aggressive malignancy of immature lymphocytes with about 15-20% of cases of T lineage (T-ALL). It is the most common type of cancer in children, but also affects adults with incidences of ∼30 new cases per 1,000,000 per year[1]. Pediatric ALL is largely curable with intensive chemotherapy, but there are significant side effects and ∼20% of patients suffer relapse. Adult ALL in contrast is largely incurable and refractory/relapsed disease is common (5-year overall survival of ∼40%)[2].

T-ALL is the result of a malignant alteration of hematopoietic progenitors in the course of through T-cell development. A relevant oncogenic pathway involved in T-cell transformation is the NOTCH1 signaling pathway with over 50% of human T-ALL carrying activating mutations of NOTCH1 gene[3],[4].

Recently, small extracellular vesicles (sEV) such as exosomes were reported to contribute to leukemic progression[5],[6]. sEV were shown to reprogram the bone-marrow microenvironment[7], dampen anti-leukemia immune response[8] and promote drug resistance[9]. sEV exert such molecular and cellular functions by transferring molecular information from cancer cells to proximal and/or to distant body districts, including pro-metastatic niche[10]. Importantly, in T-ALL a miRNA-tumor suppressor gene network drives the malignant transformation of T-cell progenitors[11],[12] and cooperates with NOTCH1-driven T-ALL[13-17]. However, the precise role of sEV and miRNA cargo in NOTCH1-driven T-ALL remains elusive. Here, we tackle this issue and present new evidences supporting a central role for EV-miRs in the progression of NOTCH1-driven T-ALL.

## Results and discussion

The molecular characteristics of sEV in T-ALL were initially explored in CUTLL1 cell line, a well-characterized human T-ALL cell line, which expresses an activated NOTCH1 and is strongly sensitive to γ-secretase inhibitors[18]. CUTLL1 cells were lentivirally transduced to constitutively express a dominant-negative form of Mastermind-like protein 1 (dnMAM) to shutdown NOTCH1 signalling, or an empty vector as a control. Indeed, HES1 expression (a target of NOTCH1) was strongly reduced under dnMAM condition (Fig. 1A, B). Next, we analyzed size distribution and quantities of sEV released from CUTLL1-CTRL and CUTLL1-dnMAM cells by nanoparticle-tracking analysis (NTA; Fig. 1C). Overall, the prevalent size of sEV matched with expected exosome size distribution (i.e., ∼30-150nm; Fig. 1C) and sEV concentration was significantly increased in dnMAM cells (Fig. 1C). We detected a total of 318 miRNAs (Fig. 1D; Table S1) by whole-miRNA expression profiling of CUTTL1 cells of which 73 also detected in sEV (i.e., Common-miRs; Fig. 1D; Table S1). Yet, hierarchical clustering analysis showed a set of highly abundant ‘EV-miRs’ comprising members of miR-17-92a cluster and paralogues (Fig. 1E). In line with previous reports, miR-19b is highly expressed in T-ALL cells and is targeted by the t(13;14)(q32;q11) translocation in T-ALL[19]. Likewise, other members of miR-17-92a clusters i.e., miR-20a and miR-92a, were found highly expressed in T-ALL and together with miR-19b were shown being capable of promoting T-ALL[11]. Expression profiling analysis of the Common-miRs in T-ALL cells and in sEV (Fig. 1D), revealed a significant and specific decreased expression of miR-17-92 cluster and paralogues upon NOTCH1 signalling inactivation (i.e., dnMAM vs. CTRL; Fig. 1F), which suggests NOTCH1 signalling modulates EV-miRs quantities in sEV. To investigate the function of these NOTCH1-dependent EV-miRs, we produced PKH26-labelled sEV enriched in miR-17-92 (aka EV_miR-17-92) by overexpressing miR-17-92 cluster in dnMAM cells (Fig. 1G-H; see methods). Internalization of EV_miR-17-92 in dnMAM cells (Fig. 1I) significantly increased the proliferation rate to a comparable level to NOTCH1-proficient CUTLL1 CTRL cells (Fig. 1J). We then treated CUTLL1-wt cells with γ-secretase inhibitor (GSI; see supplemental methods) and observed, as expected, a strong impairment of cell viability (Fig. 2A). Contrariwise, EV_miR-17-92 induced expansion of CUTLL1-wt cells (p<0.01; Student’s T-test; Fig. 2A) and, importantly, were able to rescue the GSI-induced phenotype in T-ALL cells (Fig. 2A). Similar results were obtained by using cells from two independent clones of T-ALL patient-derived xenografts (PDX) (Fig. 2A). Taken together, such results showed, for the first time, the ability of sEV_miR-17-92 to propagate molecular information among T-ALL cells which was able to restore, at least in part, a defective NOTCH1 signalling pathway.

**Fig. 1.**
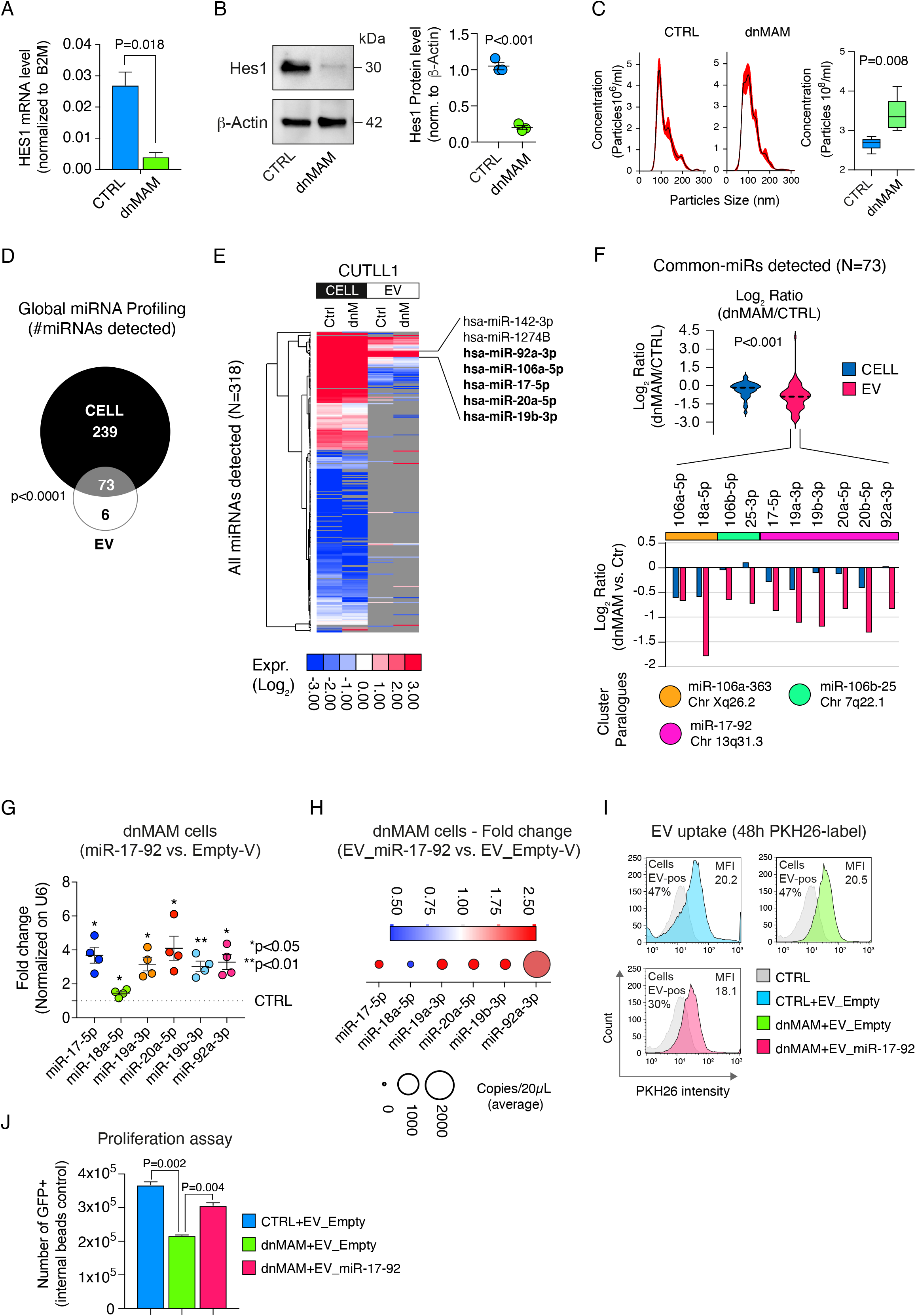
EV-miRNAs characterization and function in T-ALL model. **(A)** ddPCR analysis of HES1 mRNA expression in CUTLL1-CTRL and CUTLL1-dnMAM cells. Y-axes, Hes1 mRNA levels normalized to B2M expression. X-axes, experimental conditions. Significance analysis was performed by Student t-test. **(B)** Immunoblot analysis of HES1 expression in CUTLL1-CTRL and CUTLL1-dnMAM cells. On the right, relative quantification based on HES1 band intensities (normalized to β-actin). Significance analysis was performed by Student t-test. The experiments were performed three times **(C)** Nanoparticle-tracking-analysis of the size distribution and concentration of sEV released by CUTLL1-CTRL and CUTLL1-dnMAM cells. On the right, analysis of differential concentration of sEV in CUTLL1-CTRL and CUTLL1-dnMAM cells. Significance analysis was performed by Student t-test. **(D)** Venn diagram of EV-miRNAs detected in CUTLL1-CTRL and CUTLL1-dnMAM cells (CELL) or in their released small extracellular-vesicles (sEV). Significance analysis was performed by Fisher’s exact test. **(E)** Hierarchical clustering analysis of miRNAs detected (N=318) in CUTLL1-CTRL (Ctrl) and CUTLL1-dnMAM (dnM) cells (CELL) and/or in their released small extracellular-vesicles (sEV). On the right, most abundant miRNAs in sEV were also indicated; in bold, members of the miR-17-92 cluster. **(F)** On top, violin plots of differential expression (dnMAM vs. CTRL) of the 73 commonly detected miRNAs in CUTTL1 cells and in sEV. Bottom, bar plots of differential expression (dnMAM vs. CTRL) of the miR-17-92 cluster and paralogues. Colors are as per the legend. Significance analysis was performed by Mann–Whitney U-test. **(G)** qRT-PCR analysis of miR-17-92 cluster overexpressing CUTLL1-dnMAM cells vs. control (Empty-V) CUTLL1-dnMAM cells. Significance analysis was performed by one-sample t-test. **(H)** ddPCR analysis of miR-17-92 cluster in sEV purified from miR-17-92 cluster overexpressing vs. control (Empty-V) CUTLL1-dnMAM cells. Bubble size represents the average expression of miRNAs (copies/20µL). Colours are as per the legend. **(I)** Flow cytometry analysis of CUTTL1 (CTRL) and CUTTL1-dnMAM cells (dnMAM) conditioned with PKH26-labelled miR-17-92-enriched sEV (EV_miR-17-92) or PKH26-labelled Empty-Vector sEV (EV_Empty-V) derived from miR-17-92 overexpressing CUTTL1 cells or from CUTLL1 cells transfected with an empty vector, respectively. MFI, mean fluorescence intensity. Percentages of cells which internalized exogenous PKH26-sEV (Cells EV-pos) are also shown. **(J)** Viability of CUTTL1 (CTRL) and CUTTL1-dnMAM cells (dnMAM). Briefly, transduced GFP positive cells were FACS sorted and *in vitro* grown together with miR-17-92-enriched sEV (EV_miR-17-92) or sEV (EV_Empty-V) cultured for two days. GFP+ alive cells were measured by flow cytometry for DAPI (4′,6-diamidino-2-phenylindole) exclusion and counted by relating the cell numbers to internal fluorescent bead events (see also methods). The graph reports the result of two independent experiments. Significance analysis was performed by Student’s t-test.

**Fig. 2.**
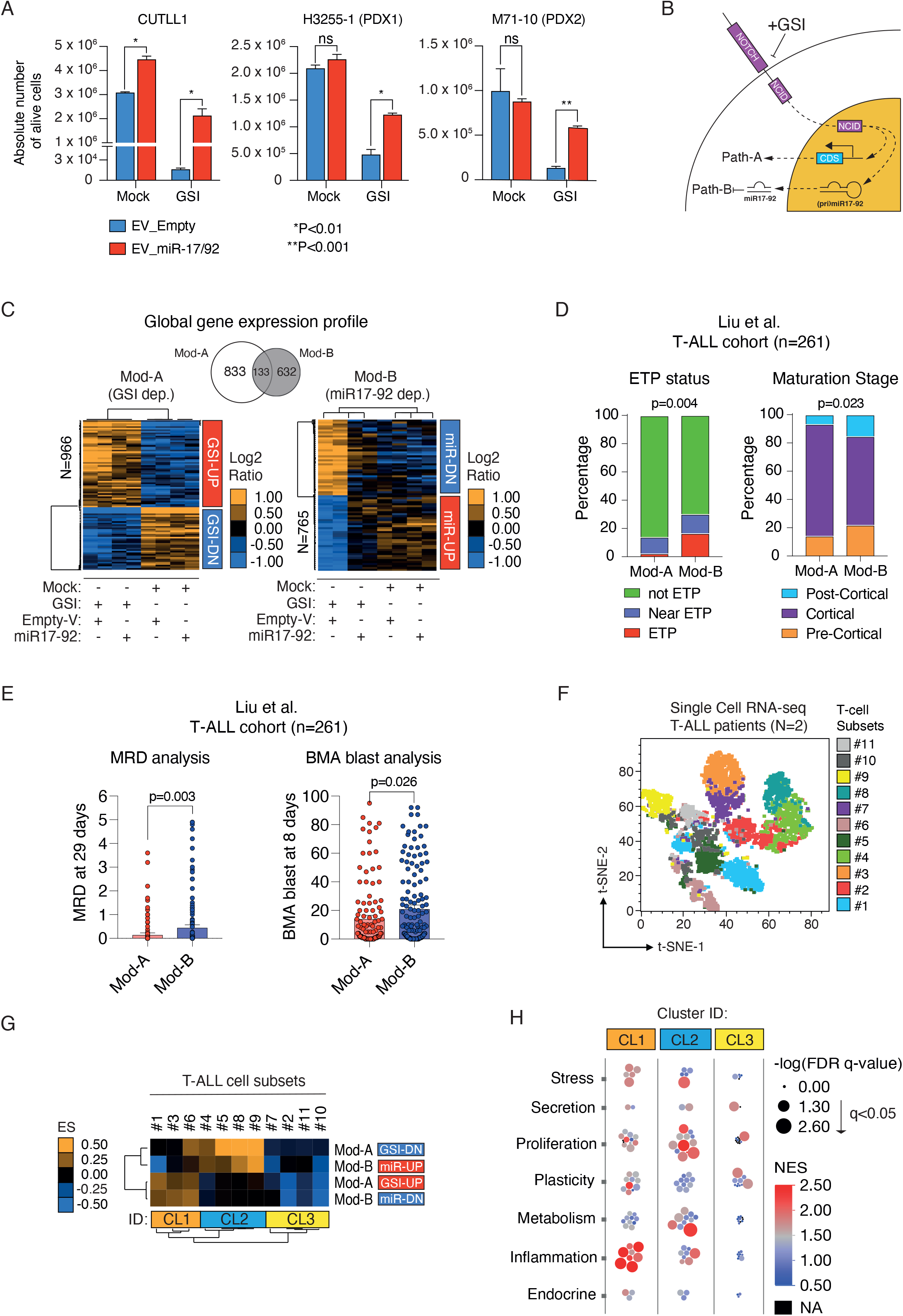
Insights in biological and molecular functions of miR-17-92 cluster in T-ALL. (**A**) Viability of CUTLL1 cells and two independent T-ALL clones of patient-derived xenografts (PDX1 and PDX2) cultured *in vitro* following lentiviral transduction with sEV from CUTTL1 cells constitutively expressing miR-17-92 cluster (EV_miR-17-92) or an empty vector (EV_Empty-V) used as control (see also methods). Transduced GFP positive cells were FACS sorted and treated with GSI (1µM, final) or DMSO as control on bare plastic for two days before flow cytometry analysis. Cell counts are related to internal beads as control. The graphs report the result of three independent experiments performed in triplicate. Significance analysis was performed by Student’s t-test. **(B)** Graphical representation of canonical (Mod-A) and of miR-17-92 modulated NOTCH1-signalling pathway (Mod-B). **(C)** Hierarchical clustering analysis of Mod-A and Mod-B gene expression profile in the various experimental conditions (i.e., +/-GSI; miR-17-92 OE or empty vector). Main clusters of genes are also indicated as GSI-UP/GSI-DOWN (Mod-A) or miR-UP/miR-DOWN (Mod-B). Colours are as per the legend. On top, Venn diagram showing Mod-A/B number of genes and relative overlapping. **(D)** Percentage distribution of ‘ETP status’ and ‘Maturation stage’ of T-ALLs in the Liu et al. cohort (n=261) stratified according to ssGSEA using Mod-A (n=123) and Mod-B (n=138) gene sets. ETP, Early T-cell Precursor. P-values were computed by chi-square test. **(E)** Box-plots show the levels of MRD (at 29 days) and BMA of blasts (at 8 days) in T-ALLs the Liu et al. cohort (n=261) stratified as in (D). MRD, Minimal Residual Disease. BMA, Bone Marrow Aspirates. **(F)** t-SNE plot of scRNAseq data on cell subsets of PDX from T-ALL patients. Colours are as per the legend. **(G)** Hierarchical clustering analysis of Enrichment Scores from GSEA using the Mod-A (GSI-UP/DOWN) and Mod-B (miR-UP/miR-DOWN) gene sets in the T-ALL cell subsets profiled by scRNAseq as in (F). **(H)** Distributions of enrichment of biological functions which were identified by GSEA using scRNA-expression profiles, in clusters of T-ALL cell subsets as in (G). The higher the size of bubbles the more significant is the enrichment of a particular biofunction. Colors of bubbles indicate the magnitude of normalized enrichment scores (NES) and are as per the legend.

Lastly, we dissected the molecular function of miR-17-92 cluster in the realm of NOTCH1-driven T-ALL. We reasoned that NOTCH1 signalling can be generalized in two main routes: Path-A) the ‘canonical’ transcriptional output of NOTCH1 intracellular domain (NCID) (Fig. 2B); Path-B) the transcriptional output controlled by NOTCH1 through miR-17-92 (Fig. 2B). High-throughput gene expression profiling of CUTLL1 cells +/- miR-17-92, and GSI/mock treated (Fig. 2C; see methods) followed by quantitative trait analysis (see methods) identified two transcriptional gene modules, i.e. Mod-A (N=966 genes) and Mod-B (N=765 genes), which differ in terms of transcriptional regulation and are both dependent to GSI treatment yet indifferent to rescued miR-17-92 expression (i.e., Mod-A; Fig. 2C; see methods), or reverted (i.e., Mod-B Fig. 2C) (Table S2). Such results confirmed our hypothesis of a bipartite NOTCH1 signalling transcriptional output (Fig. 2B). Notably, MSigDB analysis of Mod-B gene sets revealed a strong and significant enrichment (FDR q-value <0.0001) of predicted miR-17-92-targeted transcripts (Table S3; see methods) further confirming the regulatory function of the miR-17-92 cluster in Mod-B. Furthermore, IPA software (see methods) revealed that Mod-A comprised canonical NOTCH-signalling genes (e.g., NOTCH2-4, MYC, CTNNB1, GATA1-3, etc.) (Fig. S1A) while Mod-B was enriched in gene involved in proliferation (CDKN2A, CCNE1, E2F3, E2F6, RBL1), stemness (FOXM1, TCF4) and cancer (ETS1, RELA, NFE2L2) (Fig. S1B). Intriguingly, when we used Mod-A and Mod-B gene sets to stratify an external cohort of human T-ALL (i.e., the Liu et al. cohort, N=261; Table S4; [20]), we observed that Mod-B gene set hallmarks T-ALLs particularly enriched in the Early T-cell precursor (ETP) and pre-/post-cortical subtypes, with a higher post-therapeutic minimal residual disease (MRD), and blast count in the bone marrow, that are all characteristics of an adverse outcome [21-23](Fig. 2D-E; Table S5).

Next, we performed single-cell RNA sequencing of primary cells, derived from T-ALL patients (N=2), without any expansion in vivo into immunocompromised mice. Using the Phenograph algorithm [24], we identified several distinct cell subsets (n=11) (Fig. 2F). Gene set enrichment analysis (GSEA) using Mod-A and -B and hierarchical clustering analysis revealed three main clusters grouping T-ALL cell subsets which shared similar pattern of enrichment scores (ES) (Fig. 2G). In particular, CL2 contains cell subsets with coherent expression trend of both Mod-A and -B as defined in Fig. 2C, which is a hallmark of activity of NOTCH1 signalling pathway. Indeed, GSEA using Hallmark gene sets (see methods) confirmed that these CL2-cell subsets were significantly characterized by mechanisms involved in proliferation and metabolism (Fig. 2H).

Our findings shed new light on composite interactions between sEV-miRs, Notch signalling and cellular plasticity that characterize the tumor heterogeneity of T-ALL and promote relapsed/refractory cell subsets of T-cell leukemias. In this scenario, further investigations are needed to explore such mechanisms in T-ALL with the final intent of offering more efficient therapies targeting diverse oncogenic states and microenvironments that support aggressive tumor cells.

## Materials and methods

For extensive details on all methodologies see online Supplemental Material and Methods.

### Human samples

The institutional ethical committees approved this study (registration number: N91/CE), and informed consent was obtained from all patients enrolled.

### Profiling by TaqMan Human MicroRNA Arrays

Expression levels of 754 miRNAs were quantified using the TaqMan Human MicroRNA Array A+B Card Set v3.0 (Applied Biosystems, Foster City, CA).

### Genome-wide expression profiling

Gene expression profiling was performed using the GeneChip® Human Clarion S Array (Thermo Fisher Scientific) including more than 210,000 distinct probes representative of >20,000 well-annotated genes (hg19; Genome Reference Consortium Human Build 37 (GRCh37)).

### Single cell RNA-sequencing (scRNA-Seq)

Whole transcriptome analysis at single cell level was performed on FACS-sorted primary T-ALL cells using the BD Rhapsody Single-Cell Analysis System (BD, Biosciences).

## Data set availability

The normalized (U6) data for miRNA can be found in Table S1 while mRNA expression data can be accessible at NCBI GEO (GSE193482; <u>reviewer token: gjubkewovfsvdyr</u>) and SRA (PRJNA784728 for scRNA-Seq data).

## Abbreviations

T-ALL: T-cell acute lymphoblastic leukemia
sEV: small extracellular vesicles
EV-miRs: sEV contain NOTCH1-dependent microRNAs
GSI: γ-secretase inhibitors
ETP: early T-cell progenitor
dnMAM: dominant-negative form of Mastermind-like protein 1
NTA: nanoparticle-tracking analysis
PDX: patient-derived xenografts
NCDI: NOTCH1 intracellular domain
MRD: minimal residual disease
GSEA: Gene set enrichment analysis
IPA: Ingenuity Pathway Analysis
ddPCR: Droplet Digital PCR
CTRL: control
NES: normalized enrichment scores

## Declarations

## Acknowledgments

We are grateful to Chiara Di Giorgio for critically editing the manuscript and Dr. Rossella Di Paola for her help in the Illumina next generation DNA sequencing. This study was performed in accordance with the Declaration of Helsinki and was approved by the Ethics Committee of “Casa Sollievo della Sofferenza” Foundation. All authors gave their consent to publication. The gene expression data used in this study are publicly available as indicated in the Methods and Supplementary Information sections.

## Authors’ contributions

Conception and design: TC, VG, FB; Development of methodology: TC, PP, FT, VM, GR, VG, FB; Acquisition of data: TC, PP, FT; Analysis and interpretation of data: TC, PP, VG, FB; Writing, review, and/or revision of the manuscript: TC, VG, FB; Administrative, technical, or material support: PP, GR; Study supervision: VG, FB.

## Consent for publication

All authors have agreed to publish this manuscript.

