## Supplementary figures and images for "Extracellular vesicle microRNAs contribute to Notch signaling pathway in T-Cell Acute Lymphoblastic Leukemia"

### Supplementary Figure 1

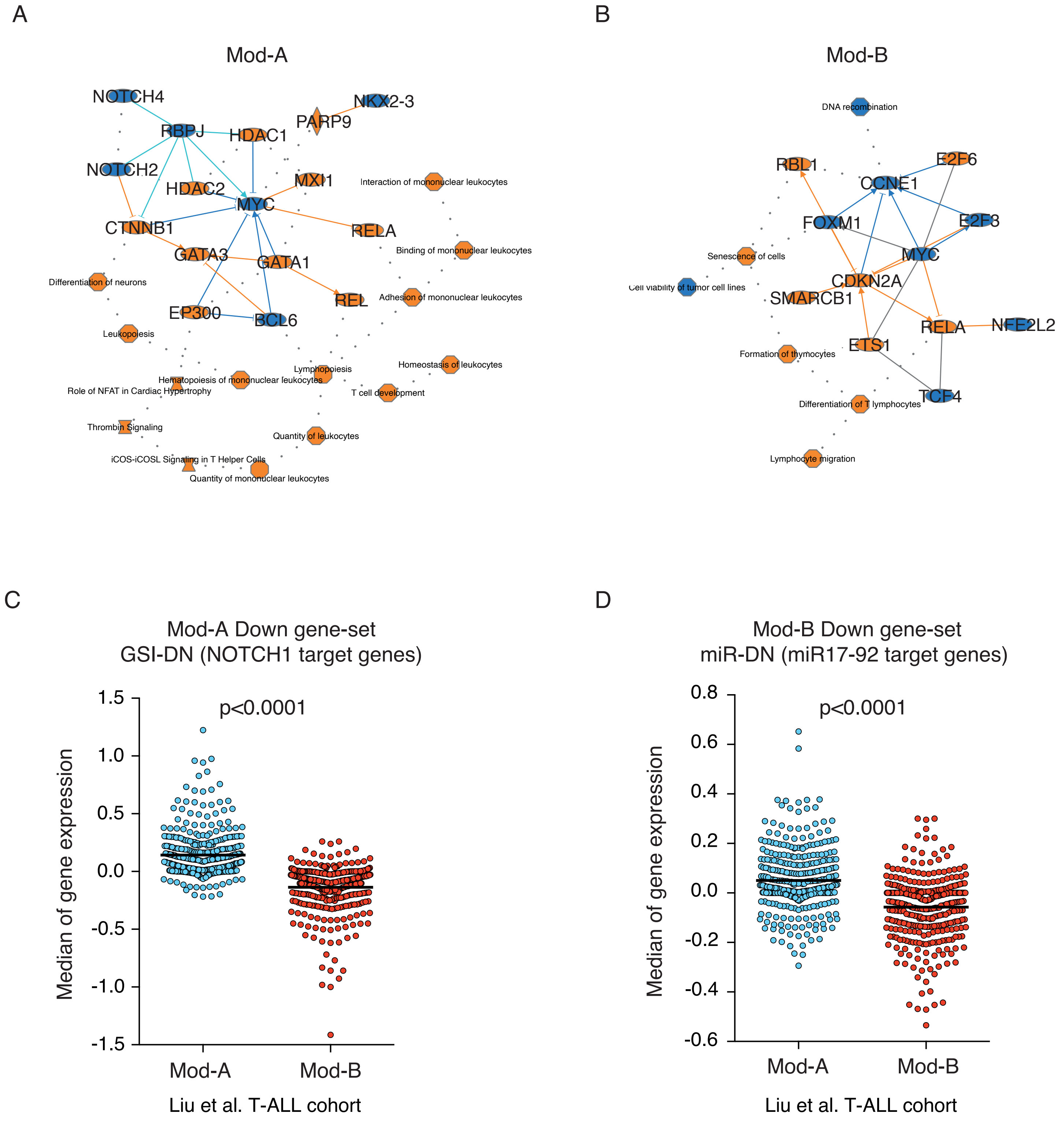
