## Supplementary Materials for "Extracellular vesicle microRNAs contribute to Notch signaling pathway in T-Cell Acute Lymphoblastic Leukemia"

**Supplemental MATERIALS and METHODS**

**PDX models and cell culture**

Patient-derived xenografts were established by injection of primary patient biopsy material into irradiated NOD-Scid/IL2Rγc^-/-^ (NSG) mice. Derivation and maintenance of M71 and H3255 PDX lines have been previously described [22]. CUTLL1 human T-ALL cell line was cultured in RPMI 1640 medium supplemented with 10% FBS, 1 mM sodium pyruvate, 2 mM L-glutamine, and antibiotics (Invitrogen). Cells were maintained at 37°C and 5% CO2 in a humidified incubator. Cells were routinely verified as free of mycoplasma contamination.

**Lentivirus vector production and transduction**

Lentivirus was produced by transient co-transfection of 293T cells with pCMVΔR8.74 and pCMV-VSV-G packaging/envelope vectors and viral transduction performed by spinfection in the presence of polybrene[22]. Virally transduced cells were sorted by flow cytometry. dnMAM, Empty-V and miR-17-92 constructs were based on pRRLsin.cPPTcts.MNDU3.PGK.GFP.WPRE backbone. Lentiviral miR-17-92 construct was derived from MSCV-mir-17-92 plasmid (cat. #64100, Addgene)[23].

**GSI treatment**

PDX cells were transduced with lentivirus encoding miR-17-92 cluster with a truncated NGFR (tNGFR) as a marker, or an empty vector as a control (CTRL), and then cultured on MS5-DL1 feeders for 5 days. Cells were subsequently sorted with FACS and then cultured without feeders in the presence of γ-secretase inhibitor GSI (1uM final) for 2 days, or DMSO as control, prior to staining for the detection of live NGFR+ by flow cytometry. Transductions were performed in triplicate.

**EV isolation and PHK26-labelling**

Small extracellular vesicles (sEV) were purified by differential centrifugation. In particular, the supernatant was collected from cells that were cultured in media containing exosome-depleted FBS (Thermo Fisher) for 72 h, and was subsequently subjected to sequential centrifugation steps at 300g for 10 min at 4 °C, and 2000g for 10 min at 4 °C. The resulting supernatant was then filtered using 0.2-μm filters to remove larger size vesicles, and a pellet was recovered at 100000g in a SW41 Ti swinging-bucket rotor after 90 min at 4 °C of ultracentrifugation (Beckman Coulter). The supernatant was aspirated and the pellet was resuspended in PBS and subsequently ultra-centrifuged at 100,000g for another 90 min. The purified EV were then resuspended in 50 μL of particle-free PBS (Gibco, Waltham, MA, USA) and used for experimental procedures.

sEV were stained using PKH26 Red Fluorescent Cell Linker Kits for General Cell Membrane Labeling according to the manufacturer’s protocol (Sigma-Aldrich). PKH26-labelled exosomes were pelleted by ultracentrifugation at 100000 ×g for 90 min at 4 °C and gently resuspended in 50 μL DPBS.

**RNA and protein extraction.**

Total RNA was extracted using RNeasy Mini Kit (Qiagen, Hilden, Germany), treated with Dnase-Rnase free (Qiagen, Hilden, Germany) and quantified by Nanodrop (Thermo Fisher Scientific). Whole protein extracts were prepared using an appropriate volume of RIPA buffer (50 mM Tris-HCl pH 7.4, 150 mM NaCl, 1% NP-40, 1% Na-deoxycholate, 1 mM EDTA, 0.1% SDS) supplemented with PhosSTOP Phosphatase Inhibitor Cocktail Tablets (Roche), and complete Mini Protease Inhibitor Cocktail Tablets (Roche).

**NTA**

sEV number and size distribution in medium samples was determined by nanoparticle tracking analysis (NTA) using a NanoSight NS300 system (Malvern Technologies, Malvern, UK). The NTA was configured with a 488 nm laser and a high sensitivity scientific CMOS camera. Samples were analysed under constant flow conditions (flow rate =30) at 25 °C and 5 × 60 s videos were captured with a camera level of 16. Data was analysed using NTA 3.4 Build 3.4.4 software with a detection threshold of 4.

**Droplet Digital PCR (ddPCR)**

First strand cDNA was generated from total RNA by reverse transcription with SuperScript III/VILO master mix (Invitrogen) including a combination of random 15-mer and anchored oligo(dT) primers. The following TaqMan probe-based assays were used: HES1 (Hs00172878_m1; FAM) and B2M (Hs99999907_m1; VIC, primer limited) (Applied Biosystems/ThermoFisher).

For miRNA expression analysis in Evs: Evs were collected from 6ml of conditioned media and Total RNA was extracted using Rneasy Mini Kit (Qiagen, Hilden, Germany), treated with Dnase-Rnase free (Qiagen, Hilden, Germany) and quantified by Nanodrop (Thermo Fisher Scientific). EV RNA was reverse-transcribed using miRScript PCR System (QIAGEN). For ddPCR reaction, 20 µL of PCR reaction containing 2µL of miScript Primer Assays (Qiagen) (Hs_miR-20a_1 (Cat.MS00003199), Hs_miR-18a_2 (Cat. MS00031514), Hs_miR-17_2 (Cat. MS00029274), Hs_miR-19a_1 (Cat. MS00003192), Hs_miR-19b_2 (Cat. MS00031584), Hs_miR-92_1 (Cat. MS00006594)), 1 µL of cDNA, 1 µL of nuclease-free water and 10 µL of 2 × EvaGreen Supermix (BioRad) were loaded into the droplet generator cartridge (BioRad). Then, 70 µL of droplet generation oil for EvaGreen (BioRad) was loaded into each of the eight oil wells and the cartridge was transferred to the QX200 droplet generator (BioRad). The droplets (40 µL) were, then, transferred to a 96-well PCR plate (BioRad) and the plate was heat-sealed with specific aluminum foil (BioRad). Thermal cycling conditions for EvaGreen assays were as follows: 95 °C for 5 min, 40 amplification cycles of 94 °C for 15 s, 55 °C for 30 s and 70 °C for 34 s, and three final steps at 4 °C for 5 min, 90 °C for 5 min, and a 4 °C indefinite hold to enhance dye stabilization. Finally, the dropletized PCR was read by the Q×200 Droplet Reader and the Poisson-corrected determination of template concentration was calculated using QuantaSoft™ Analysis Pro Software (v1.0, Bio-Rad). For quantification, a minimum of 10,000 acceptable droplets per 20 µL reaction was used, followed by manual selection of positive and negative droplet populations.

**Quantitative Real Time-PCR (qRT-PCR)**

miRNA expression analysis was performed using 1µg of total RNA was reverse transcribed using miRScript PCR System and analyzed by qRT-PCR using the miScript SYBR Green PCR Kit. The miScript Primer Assays (Qiagen) were used following manufacturers’ instructions and in particular: Hs_miR-20a_1 (Cat.MS00003199), Hs_miR-18a_2 (Cat. MS00031514), Hs_miR-17_2 (Cat. MS00029274), Hs_miR-19a_1 (Cat. MS00003192), Hs_miR-19b_2 (Cat. MS00031584), Hs_miR-92_1 (Cat. MS00006594). For qRT-PCR data normalization we used the Hs_RNU6-2_1 (Cat. MS00033740) miScript Primer Assay (Qiagen) which quantifies U6 expression level and data were processed. qRT-PCR experiments were performed using a QuantStudio 12K Flex Real-Time PCR System (ThermoFisher Scientific) with manufacturer’s recommended cycling conditions (Qiagen). Log2 ratios of miR-expression levels were obtained using the 2-ΔΔCT method.

**Western Blotting**

Cell pellets were incubated for 30 minutes on ice. Lysates were cleared by centrifugation at 17000 × g for 30 minutes at 4°C. Protein concentration was determined by Pierce BCA Protein Assay kit (ThermoFisher Scientific) according to manufacturer’s instructions. 30µ of whole protein lysate were loaded onto Mini-protean TGX gels (Bio-Rad, Hercules, CA, USA) and then transferred on a polyvinylidene fluoride (PVDF) membrane (Bio-Rad) using Trans-blot Turbo (Bio-Rad). Membranes were blocked for 1 h at room temperature (RT) with either 3% BSA/0.1% Tween-20/TBS/(w/v/v) or with 5% non-fat dry milk/0.1% Tween-20/TBS (w/v/v), and then incubated overnight (ON) at 4 °C with primary antibodies (anti-HES1, ab71559, Abcam; anti-𝛽-actin, sc-47778, Santa Cruz). Membranes were then incubated for 1h at RT with goat anti-mouse IgG or goat anti-rabbit IgG secondary antibodies (Promega, Madison, WI, USA). Proteins were visualized using ClarityTM or Clarity MaxTM Western ECL substrates (Promega, Madison, WI, USA). Images were acquired using the ChemiDoc™ Imaging system (Bio-Rad). Protein levels were quantified using the Image Lab software (version 5.2.1 build 11, Bio-Rad). Experiments were repeated at least three times.

**Profiling by TaqMan Human MicroRNA Arrays**

Briefly, 3 μL of total RNA (EV profiling) or 10ng of total RNA (intracellular profiling) were reverse-transcribed into cDNA by the TaqMan MicroRNA Reverse Transcription Kit and Megaplex RT set pool A and B v3.0 (Applied Biosystems). The resulting cDNAs (5 μL) was then pre-amplified for 14 cycles using the corresponding Megaplex™ PreAmp Primers and TaqMan PreAmp MasterMix. Real-time quantitative PCR (RT-qPCR) was carried out with 18 μL of 1 in 4 diluted cDNA template was diluted and mixed with TaqMan Universal PCR Master Mix, and loaded into each of the eight fill ports on the TaqMan Array. TaqMan MicroRNA Arrays were then run using the QuantStudio 12K Flex Real-Time PCR System (ThermoFisher Scientific). All reactions were performed according to standard manufacturers' protocols.

**Genome-wide expression profiling**

RNA samples have been amplified, fragmented, and labeled for array hybridization according to the manufacturer’s instruction. Samples were then hybridized ON, washed, stained, and scanned using the GeneChip Hybridization Oven 640, Fluidic Station 450 and Scanner 3000 7G (Thermo Fisher Scientific) to generate the raw data files (.CEL files). Quality control and normalization of CEL files were performed using Transcriptome Analysis Console (TAC) software v4.0 (Thermo Fisher Scientific) by performing “Gene level SST-RMA” summarization method with human genome version hg38. The analyses were performed in two independent biological experiments and gene expression data were Log2 transformed before analyses.

**Flow Cytometry**

An anti-hCD271 (eBioscience, ThermoFisher) was used to detect the lentiviral NGFR. We performed FACS analysis and sorting on FACS Calibur, Canto2, and Aria2 cytometers (Becton Dickinson) and MoFlo Astrios cell sorter (Beckman Coulter). We analyzed flow cytometry data using FlowJo software (Becton Dickinson). The proliferation assay was performed by counting absolute cell numbers with the AccuCheck counting beads (cat. PCB100, ThermoFisher) according to the manufacturer’s instructions. Cell viability was measured by propidium iodide exclusion.

Leukemia cells were co-incubated with 2x10^8^ PKH26-labeled EV, derived from CUTLL1 empty-vector or miR-17-92 cluster, for 48h at 37C in a media containing EV-depleted FBS and washed repeatedly to remove unbound EV. Cellular internalization of PKH26-positive EV was measured by MoFlo Astrios cell sorter (Beckman Coulter).

**Single cell RNA-sequencing (scRNA-Seq)**

Leukemia cells from cryopreserved bone marrow samples of T-ALL patients were isolated as CD45+CD3-CD99+CD7+ cell fraction by FACS-sorting as previously reported[24]. Isolated cells from each sample were labeled with the BD Single-Cell Multiplexing Kit (cat. 633781, BD Biosciences) for 1 hour following the manufacturer’s protocol. The cell viability and concentration were determined with the BD Rhapsody Scanner system after staining with viability dyes, Calcein AM (1:200 dilution; cat. #C1430, ThermoFisher) and DRAQ7^TM^ (1:200 dilution; cat. #564904, BD Biosciences), and incubation for 5 min at 37°C. Cells were counted using the Improved Neubauer Hemocytometer (INCYTO). Afterward leukemia cells for each sample at d0 and d30 were pooled equally in 650ml cold BD Sample Buffer and three BD Rhapsody cartridges were loaded with 10,000 pooled cells for single cell separation. Single cells were isolated using Single-Cell Capture and cDNA Synthesis with the BD Rhapsody Express Single-Cell Analysis System according to the manufacturer’s recommendations (BD Biosciences). Based on the number of viable cells revealed and captured on the beads, the final resuspension volume was calculated to subsample and sequence about 4,000 cells. Whole transcriptome and Sample Tag amplification was performed with the BD Rhapsody Whole Transcriptome Amplification Kit (cat. #633774) following the manufacturer’s instructions. Unwanted PCR products and other small molecules were excluded performing a side cleanup using the AMPure XP Beckman magnetic beads (cat. #A63880, Beckman Coulter). DNA quantity and quality control were performed using the Qubit^TM^ dsDNA HS Assay Kit (cat. # Q32851, ThermoFisher Scientific) and the electrophoresis system Agilent 2200 TapeStation, cartridge (cat. #5067-5584). Sequencing was performed in paired-end mode (2*75 cycles) on NextSeq 500 System (Illumina) with the NextSeq 500/550 High Output Kit v2.5 (150 Cycles) chemistry to reach a depth of 75,000 reads for WTA and 500 reads for SMK per cell for a total of 80,000 reads per cell on average. Sequencing data were processed on the Seven Bridges Platform (<https://www.sevenbridges.com/>) for sample demultiplexing and generation of expression sparse matrixes. Briefly, a quality sequencing step was performed to filter out reads with a Phred quality score below 20, reads with a single type of nucleotide and short reads (less than 60 bases for R1 and less then 42 for R2). UMI counting was done including sequencing reads correction, recursive substation error correction (RSEC) and distribution-based error correction (DBEC). Accepted reads were aligned to the reference genome (GRCh38) for identification and quantification and subsequently the count genes matrix was generated. Highly dimensional Sc-RNA-Seq data were analyzed using the SeqGeq software (Becton Dickinson) for visualization, clustering and differential expression analysis. Only cells with mitochondrial read rate ≤ 30%, detectable genes ≥ 200 and genes expressed in 10 or more cells were considered passing the QC and further analyzed using functions provided with the Seurat library. In the end, we counted 6,322 total cells. Data were log normalized regressing out both the number of counts and percentage of reads aligning to mitochondrial genes. Clusters were allocated to cell populations based on gene markers using uncentered correlation and centroid linkage.

**Bioinformatics analysis**

*Quantitative Traits analysis.* Quantitative Traits (QT) consist of “measurable phenotypes that depend on the cumulative actions of many genes and the environment (www.nature.com/subjects/quantitative-trait). The BRB ArrayTools_v4.6 software (https://brb.nci.nih.gov/BRB-ArrayTools/) was used to identify Mod-A and Mod-B gene sets using the Quantitative Trait Analysis tool (Pearson correlation, cutoff ρ>0.8; FDR<10% based on 1000 random permutations for multivariate analysis; confidence level of false discovery rate assessment: 80 %). Heatmaps and clusters were generated by using Cluster 3.0 for Mac OS X (C Clustering Library 1.56) and Java TreeView version 1.1.6r4 (using Euclidean distance and centroid linkage for Figure 1E, or uncentered correlation and centroid linkage for Figure 2C, G). Analysis of the enriched biological themes in Mod-A and Mod-B gene sets (Figure 2C; Table S2) was performed either by using Ingenuity Pathway Analysis (IPA; Qiagen) with “Graphical Summary function” (Supplemental Figure S1A-B) and by using Molecular Signature Database (MSigDB; www.gsea-msigdb.org/gsea/msigdb/index.jsp) with the MIR:MIR_Legacy set collection (N=221 gene sets) and q-value (FDR) <0.05 as a cutoff for statistical significance of overlapping (Table S3). Gene Set Enrichment Analysis (https://www.gsea-msigdb.org/gsea/index.jsp) of scRNA-seq data was performed using Mod-A and Mod-B gene sets (N=4; Figure 2G; Table S2) or HALLMARK gene sets (N=50; <http://www.gsea-msigdb.org/gsea/msigdb/collections.jsp>; Figure 2H) and with 1000 random gene sets permutation. Genes were annotated to “Human_Gene_Symbol_with_Remapping_MSigDB.v7.2.chip”. Bubble plots analysis (Figure 2H) was performed using JMP 15 software (SAS Institute Inc., Cary, NC, USA) while all other plots were generated using GraphPad Prism 7 (GraphPad Software Inc., San Diego, CA, USA). All statistical analyses were performed using GraphPad Prism 7 (GraphPad Software Inc.) and JMP 15 (SAS Institute Inc.). The type of statistical test used in the various analyses is indicated in the relative figure legend. A p-value less than 0.05 was considered for statistical significance.

*Gene expression analysis of patients.* We retrieved gene expression (RNAseq) data and clinical information from the T-ALL dataset published by Liu et al. (N=264)[17]. FPKM reads were median centered before analysis and clinical data were merged into a single file. ssGSEA (single sample Gene Set Enrichment Analysis[25]) was applied to normalized FPKM using Mod-A and Mod-B gene sets (N=4; Figure 2G; Table S2). To obtain a unique score to classify each T-ALL samples into Mod-A or Mod-B enriched, we ranked the ssGSEA scores of each gene set (the highest ssGSEA score corresponds the lowest rank) and we made additions of UP and DOWN scores for each patient. If the summed rank was lower in Mod-A compared to Mod-B, then the patient was assigned to ‘Mod-A’ class otherwise to ‘Mod-B’ class. Three out of 264 patients were excluded because they had an identical score for ‘Mod-A’ or ‘Mod-B’ class (Table S4).

**Supplemental Figure Legends**

**Figure S1. (A-B)** Ingenuity Pathway Analysis of Mod-A and Mod-B genes modules.

**Supplemental Table Legends**

**Table S1. miRNA normalized (U6) expression data.** Normalized microRNA expression data in CUTLL1 cells at intracellular level (CELL) or in extracellular vesicles (EV), in dnMAM or CTRL conditions. Data are expressed as dCT (i.e., the difference of CT of the miRNA of interest and of U6 gene used as housekeeping), or as -ddCT (i.e., Log 2 Ratios, dnMAM vs. Ctrl).

**Table S2.** Mod-A and Mod-B genes. Mod-A_UP, genes upregulated upon GSI treatment. Mod-A_DOWN, gene downregulated upon GSI treatment. Mod-B_Up, genes upregulated upon miR17-92 OE. Mod-B_DOWN, genes downregulated upon miR17-92 OE.

**Table S3. MSigDB analysis of Mod-B genes.** Results of analysis of overlapping of Mod-B gene set with C3 (miRNA targets) gene sets available in the Molecular Signatures Database (MSigDB). # Genes in Overlap (k), number of genes of Mod-B gene set overlapping with the indicated miRNA targets gene set (‘Gene set name’); k/K, fraction of Mod-B genes (k) overlapping with the genes in the MSigDB gene set (K); p-value, significance of overlapping (hypergeometric distribution); FDR q-value, false discovery rate analog of hypergeometric p-value after correction for multiple hypothesis testing according to Benjamini and Hochberg.

**Table S4.** ssGSEA classification of Liu et al. T-ALL cohort using Mod-A and Mod-B gene sets (N=261 patients)**.**

**Table S5.** Clinicopathologic characteristics of the Liu et al. T-ALL cohort (ALL) and of patients classified as Mod-A or Mod-B using ssGSEA.
